# First Osteosarcoma Reported from a New World Elapid Snake and Review of Reptilian Bony Tumors

**DOI:** 10.1101/461202

**Authors:** Alexander S. Hall, Justin L. Jacobs, Eric N. Smith

## Abstract

Cancer chiefly occurs in vertebrates. Rare in amphibians, and perhaps common in reptiles, various neoplasms and malignant cancers have been reported with erratic frequency by museums, paleontologists, veterinarians, and pet hobbyists. Unsurprisingly, most herpetofaunal diversity has never been systematically surveyed for the presence of neoplasms owing to the extreme rarity or obscurity of many species. Museum collections can fill these gaps in knowledge, especially when researchers use non-destructive techniques. In this study, we used X-ray computed tomography to discover and characterize an osteosarcoma of the spine in a rare South American coralsnake, *Micrurus ancoralis.* Two spinal vertebrae were completely fused and adjacent vertebrae showed evidence of corruption. The fused vertebrae contained a hollow inner network thought to be vascular tissue. We also review previous reports of tumors in the Elapidae and all bony tumors in non-avian reptiles. The rarely reported technique of X-ray CT for tumor discovery could greatly improve our understanding of the species diversity and perhaps underlying causes of neoplasia.

---

Neoplasms, abnormal cell growths, can grow to large sizes and are often called tumors or cancers (Cooper, 1992; Hernadez-Divers and Garner, 2003). Tumors can be benign or malignant, and chiefly occur in vertebrates (Capasso, 2005). Neoplasms are reported across the entire spectrum of vertebrates (Schlumberger and Lucke, 1948; Stolk, 1958; Capasso, 2005; Rothschild et al., 2012). Amphibians exhibit neoplasms rarely (Capasso, 2005), though there are a few documented cases (Schlumberger and Lucke, 1948; Hill, 1954, 1955; Effron et al., 1977; Hruban, et al. 1992; Rothschild et al., 2012). In contrast, extant reptiles exhibit a diverse range of tumors (Machotka, 1984), and nearly all tumors found in mammals can be found in reptiles plus a few that can’t be found in mammals (Effron et al., 1977; Garner et al., 2004; Suedmeyer, 1995; Mauldin and Done, 2006; Rothschild et al., 2012). Our knowledge of reptilian neoplasms is primarily gleaned from systematic reviews from zoos (Hill, 1952-1955, 1957; Cowan, 1968; Ippen, 1972; Effron et al., 1977; Ramsay and Fowler, 1992; Ramsay and Lowenstein, 1996; Catão-Dias and Nichols, 1999; Sykes and Trupkiewicz, 2006), though important contributions have come from the pet trade and veterinary practice (Garner et al., 2004; Mader, 2005; Simard 2016; Christman et al., 2017), private collections (Frye, 1994), and museum collections (Harshbarger, 1969; Machotka and Whitney, 1980).

The frequency of tumors in vertebrates is unclear. A clear historical bias exists in the literature reflecting improved screening and expectations for what animals experience neoplastic growths, and only obvious cases were then reported (e.g., Ohlmacher, 1898; Kalin, 1937). For instance, Peyron (1939) dissected 4,000 snakes and reported no tumors. Similarly, Bergman (1941) noted only one tumor in over 2,200 snakes. The occurrence of specific tumors also changes through time. As one example, Dawe and Harshbarger (1969) explain that reptiles lack leukemias, leukoses, or reticular neoplasms, yet Frye and Carney (1973) report a lymphatic leukemia in a boa constrictor only a few years later. In 1948, Schlumberger and Lucke presented an impressive review of fish tumors and commented on the state of tumor descriptions in vertebrates. They lamented that previous investigators did a rather poor job at surveying neoplasms in reptiles and this led to a misconception that reptilian neoplasms were rare or absent. Scrutinizing specimens with greater rigor clearly yielded a higher incidence of neoplasia than previously reported. For example, in the same paper they reported over 1,200 examples of renal adenocarcinoma in the frog *Rana pipiens* in a 15 year period (Schlumberger and Lucke, 1948).

One early effort to systematically report on vertebrate neoplasms was the annual report from the Zoological Society of London prosector (e.g., Hill 1952-1955, 1957). Here, all London Zoo causes of death were reported. These annual reports included unexpected tumors, thus raising the profile of neoplasms in non-mammals. Recognizing the paucity of data and importance of comparative information of neoplasms in animals, in 1969 the Smithsonian Institution established the Registry of Tumors in Lower Vertebrates (RTLA) to allow for a common repository for and study of such specimens (Harshbarger, 1969). The Armed Forces Institute of Pathology and the RTLA were critical in establishing the fact, now widely accepted, that tumors are widespread in vertebrates and are found in nearly all tissue types (Machotka and Whitney, 1980; Frye, 1994; Hernandez-Divers and Garner, 2003).

The question of neoplasm incidence or rate in vertebrates is still the subject of research and varies depending on the context of what incidence rate is being calculated. Among birds and mammals, neoplasms are apparently rare in wild populations, but far more likely in captive animals (Bergman, 1941; Petrak and Gilmore, 1982; Jubb and Kennedy, 1970; Capasso, 2005). Similarly, as previously reported, Bergman (1941) found only one tumor in 2,200 free-ranging snakes. Effron et al. (1977) provided one of the first modern, authoritative estimates of neoplasia in vertebrates from over 10,000 necropsies at the Zoological Society of San Diego. They reported a 3% rate of neoplasia in mammals and 2% in reptiles and birds. None of the amphibians dissected exhibited neoplasia. A more recent report from an exotic species pathology service hypothesized that, among reptiles, neoplasia may be most common in snakes (Garner et al., 2004). These authors found that neoplasia in snakes are most prevalent in crotalids, followed by viperids, and then boids (including elapids). Simard (2016) recently reported 0.9% of 3995 reptiles available to Ghent University had some kind of neoplasm. Most cases were based on assessment of ante-mortem diagnostics; thus this estimate may be low since only very obvious neoplasms were reported. As reptiles are increasingly owned as pets (Collis and Fenili, 2011), systematic reports from the pet trade and associated veterinary profession will be crucial for expanding our knowledge of the incidence of tumors, at least in captive animals. However, this trend of neoplasm reporting will likely generate a bias towards those animals most commonly kept as pets. Thus, natural history collections will remain useful supplements to study neoplasia, especially in rare and ‘uncharismatic’ species not found in the pet trade (Keck et al., 2011).

In this report, we describe a bony, vascular tumor (osteosarcoma) of the new world elapid coralsnake, *Micrurus ancoralis* (Jan, 1872). Our discovery was made as part of a larger, systematic study of vertebral morphology in elapid snakes using computed tomography (CT). Prior to this report, we are unaware of any published neoplasms in new world elapid snakes (see Schmidt, 1936; Slowinsky, 1995 for phylogeny). Furthermore, this may be the first time that CT has been used as an exploratory tool for finding and describing a tumor in a vertebrate museum specimen. Rosenberg et al. (1995), demonstrated difficult cases for describing osteosarcomas using plain radiographs and supplements known osteopathic cases with tomograms from CT. Schumacher and Toal (2001) enumerate radiography and ultrasonography techniques for reptile veterinary medicine but discount CT as a routine diagnostic tool. Banzato et al (2013a, b) review veterinary imaging for snakes and lizards and include traditional CT and contrast-enhanced in their review. Cole et al. (2014) used micro CT to describe osteosarcomas in mice injected with an osteosarcoma cell line, confirming that CT can accurately describe osteosarcoma without dissection. We also review known elapid tumors and bony tumors in reptiles.

## Methods & Materials

### Computed Tomography

The formalin-fixed specimen was CT scanned at UT Arlingon’s Shimadzu Center for Environmental Forensics and Material Science using a Shimadzu inspeXio SMX-100CT (Shimadzu, Kyoto, Japan). The snake was scanned twice using similar parameters, but with different focal areas. Energy settings were 40kV and 40μA. All slices averaged 6 frames on each of 4 rotations for a total of 1200 views per full rotation. The first, focal scan consisted of 876 slices with 9 μm isotropic voxels. A second, multi-loop section with more vertebrae in focus consisted of 1697 slices with 11 μm isotropic voxels. Each scan lasted about 15 min.

Raw X-ray data were reconstructed using Shimadzu’s inspeXio software and exported as 1024 x 1024 16-bit TIFF. Each stack of 16-bit TIFF images was rotated and cropped in ImageJ and imported into Thermo Scientific™ Avizo™ Software 9.5 (Thermo Fisher Scientific, Waltham, MA, US). As a volume, each data set was resampled to align better with the X axis, cropped, filtered using ring-artifact correction, passed through a non-local means filter, and then filtered using unsharp masking. This combination of filters retained fine features and smoothed the majority of artifacts found in all computed tomography data. Multiloop scans in cone beam CT inherently experience distortion at the periphery of each component scan, and this is slightly noticeable in the final, filtered data. Vertebrae were segmented in Avizo following Hall et al. (2018). In the multi-loop scan, pores in the fused vertebrae were segmented out using a combination of thresholding and manual editing. The raw CT data, videos of each data set (Vids. S1-S3), and label data are available on a MorphoSource project: http://www.morphosource.ors/Detail/ProiectDetail/Show/proiectid/625. Additionally, raw CT data are archived at the University of Texas at Arlington Amphibian and Reptile Diversity Research Center digital image collection.

### Literature Reviews

We sought to generate two reviews of neoplasms in snakes: one specific to the snake family Elapidae, and another for all bony tumors in non-avian reptiles. We compiled a list of elapid neoplasms from a thorough literature search using relevant search terms in English, Spanish, German, French, and Dutch. This included searches with combinations of all genera of Elapid snake (reconciled between Campbell and Lamar [2004] and Uetz [2018]), the terms “elapid,” “elapinae,” or “Elapidae,” and “tumor,” “cancer,” “neoplasm,” or “sarcoma” etc. Similarly, we searched for bony tumors – defined as neoplasms of osteoids, periosteum, skeletal cartilage, and adjacent supporting tissues in the case of metastasis – in combination with common and scientific names for orders and family names of reptiles, per Uetz (2018). In each review, we examined all literature that cited relevant articles and also the references cited within. We did this regardless of whether these secondary articles could be found in the searches laid out above. Given the historical utility of the RTLA (e.g., Harshbarger, 1969), we contacted their current curatorial home to check for any unreported, relevant neoplasms for these reviews. For internal consistency, we updated common and scientific names throughout this report to match The Reptile Database (Uetz 2018).

## Results

### Computed Tomography

Two cervical vertebrae were fused together in bone and vasculature (Fig. 1C, D; Vids. S1, S3). The dorsal region of these two vertebrae was porous and resembled trabecular bone. Fusion of the vertebrae increased from the ventral to dorsal aspect. The surface and interior of the neural arch was entirely fused between the two vertebrae and its surface was porous. In contrast, the surface of cervical vertebrae elsewhere in the same specimen were smooth and not porous (Fig. 1A, B, Vid. S2). The condyle of the anterior fused vertebra and the adjacent cotyle were fused with some porosity. Unaffected vertebrae exhibit a uniform gap. Similarly, the postzygaphyseal and prezygapophyseal articular facets of the affected vertebrae were fused with some porosity. Unaffected vertebrae had a uniform gap (Fig. 1A, Vids. S1, S2). The anterior aspect of the neural spine of the posteriorly adjacent vertebra was split in a Y-shape and porous. The neural spines of the two anteriorly adjacent vertebrae were porous, dorsolaterally blunted, and enlarged on the anterior aspect. The surface of the inside of the neural canal in fused vertebrae was rougher and more porous than unaffected vertebrae. The hypaphosyes of fused vertebrae were not visibly different compared to unaffected vertebrae. Canals, thought to be blood vessels, ran chaotically through each affected vertebra and seamlessly crossed the two vertebrae (Fig. 1D, Vid. S3). By contrast, blood vessels in unaffected bones only ran through the center of each lateral vertebral process, condyle, cotyle, and hypaphosis (Fig. 1B).

**Figure 1.**
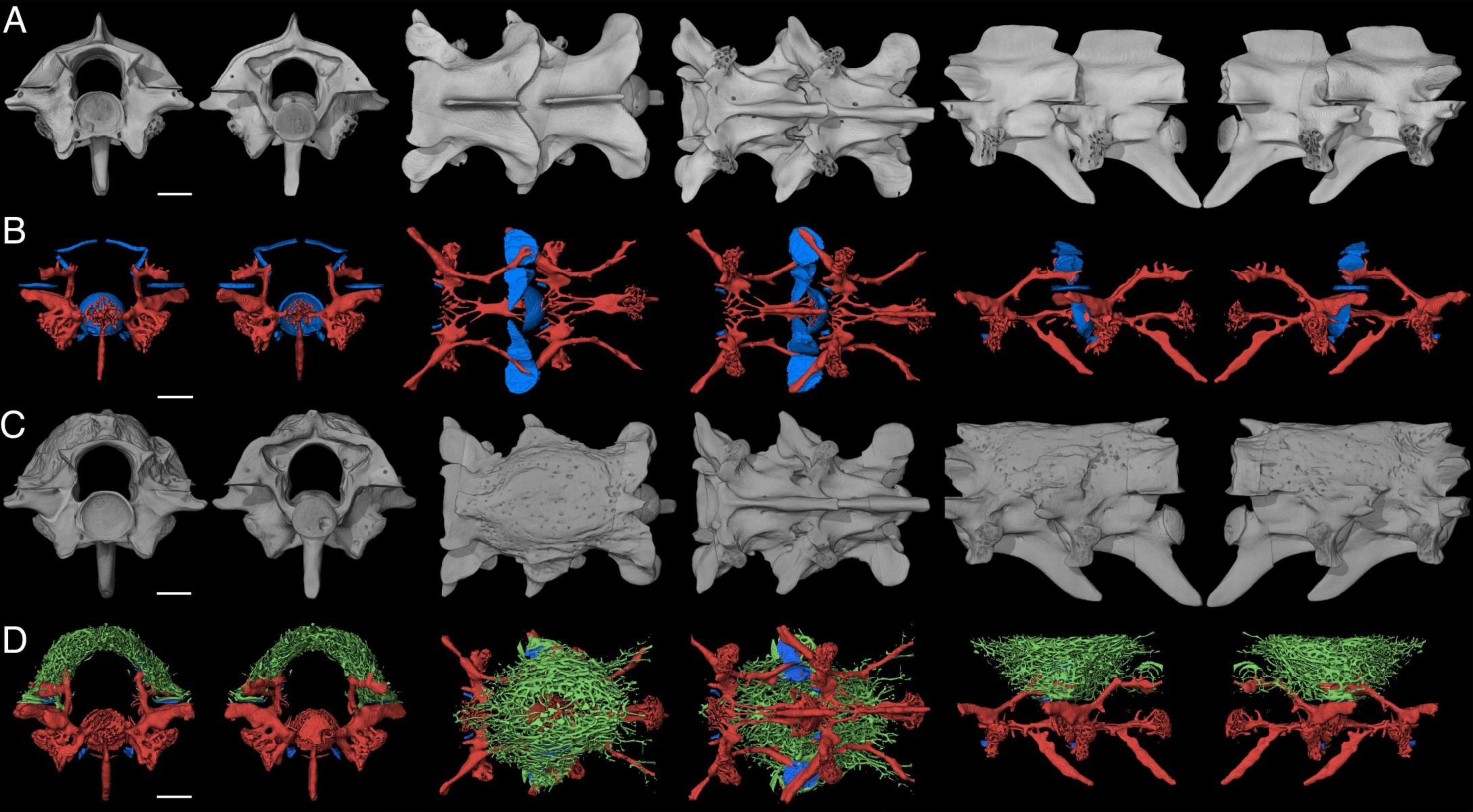
Vertebral osteosarcoma in a new world elapid snake, Micrurus ancoralis (UTA R-55945). Pairs of vertebrae from unaffected (A, B) and pathologic (C, D) regions of the spine as shown with micro-computed tomography. Blood vessels (in red) and ligament and cartilage (in blue) are shown (B and D) for each pair. Chaotic vasculature is shown in green for pathologic vertebrae (D). From left to right, views are cranial, caudal, dorsal, ventral, and both lateral angles. Scale bars are 1 mm.

### Literature Reviews

We found 38 cases of neoplasia in the snake family Elapidae, all from old world species (Table 1). Of 13 cases when sex was reported, four were male and nine were female. 23 (60%) of cases reported were from the genus *Naja,* with the second most cases from *Aspidelaps* (four; 10.5%) and then third most from *Ophiophagus* (three; 7.9%). Kidney-related neoplasms were the most reported (nine; 23.7%; Keck et al. 2011). Despite the limited number of reported elapid neoplasms, reports covered many cell types and locations (Table 1), including three bony tumors in *Naja* species. Our review of bony tumors in reptiles revealed 58 cases (Table 2). This included 49 squamates, 5 turtles, and 4 crocodylians. Six of ten cases with known sex were female and four were male. We found several varieties of bony neoplasms, including osteoma (bone cell neoplasm), osteosarcoma (malignant bone cell tumor), osteochondroma (bony, cartilage covered exotoses), chondroma (cartilage cell neoplasm), and chondrosarcoma (malignant cartilage cell tumor). *Varanus* and *Pantherophis* each included eight cases of bony tumors (16.3% of squamates). We note very few bony tumors of the head or face: only one report of osteoma of the skull from a male *Caiman crocodilus* (Kalin 1937).

**Table 1.** Neoplasms found in the snake family Elapidae.

| Scientific Name | Common Name | Neoplasm | Reference |
| --- | --- | --- | --- |
| *Acanthophis antarcticus* ♀ | Common Death Adder | Lymphosarcoma, leukemic | Effron et al., 1977 |
| *Aspidelaps l. lubricus* | Cape coral snake | Tubular carcinoma (kidney) | Keck et al., 2011<br>(unpublished) |
| *Aspidelaps l. lubricus* ♀ | Cape Coral Snake | Renal adenocarcinoma | Keck et al., 2011 |
| *Aspidelaps l. lubricus* ♀ | Cape coral snake | Renal adenocarcinoma | Keck et al., 2011 |
| *Aspidelaps l. lubricus* ♂ | Cape Coral Snake | Renal adenocarcinoma | Keck et al., 2011 |
| *Acanthophis madagascarensis*<br>(92-SnN-75)* | ?? | Squamous cell carcinoma | Frye, 1994 |
| *Dendroaspis polylepis* | Black Mamba | Biliary adenocarcinoma with sarcomatous<br>desmoplasia | Ramsay et al., 1996 |
| *Naja atra* | Chinese Cobra | Carcinoma (pancreas?) | Machotka and Whitney,<br>1980 |
| *Naja haje* (87-B-51) | Egyptian cobra | Papillary adenoma | Frye, 1994 |
| *Naja melanoleuca* | Black-and-white Cobra | Venom gland carcinoma | Hill, 1952 |
| *Naja melanoleuca* | Black Cobra | Osteochondrosarcoma (spinal column) | Wadsworth, 1954a, b |
| *Naja melanoleuca* (?) | “African Black Cobra” | Osteochondroma (dorsal aspect of spinal<br>column) | Wadsworth, 1954b |
| *Naja melanoleuca* ♂ | Black Cobra | Ductular adenocarcinoma (pancreas) | Sykes and Trupkiewicz,<br>2006 |
| *Naja naja* | Indian Cobra | Biliary adenoma | Ramsay et al., 1996 |
| *Naja naja* | Indian Cobra | Renal adenoma | Ramsay et al., 1996 |
| *Naja naja* | Indian Cobra | Adrenal adenoma | Ramsay et al., 1996 |
| *Naja naja* | Indian Cobra | Bile duct adenoma | Cowan, 1968 |
| *Naja naja* | Indian Cobra | Ganulosa cell tumor | Ramsay et al., 1996 |
| *Naja naja* | Indian Cobra | Pancreatic carcinoma | ?? |
| *Naja naja* | Indian Cobra | Granuloma (head and neck) | Wadsworth, 1956 |
| *Naja naja* (80-68452) | Indian Cobra | Seminoma | Frye, 1994 |
| *Naja naja* (RTLA 560) | Indian Cobra | Leiomyosarcoma | Machotka and Whitney, 1980 |
| *Naja naja* ♀ | Indian Cobra | Hepatocellular carcinoma (liver) | Catão-Dias and Nichols, 1999 |
| *Naja naja* ♀ | Indian Cobra | Hepatocellular carcinoma (liver) | Catão-Dias and Nichols, 1999 |
| *Naja nigricollis* (90-SB-63) | Black-necked Spitting Cobra | Renal, Tubular adenoma | Frye, 1994 |
| *Naja nigricollis* ♀ | Black-necked Spitting Cobra | Lymphosarcoma | Effron et al., 1977 |
| *Naja nigricollis* ♂ | Black-necked Spitting Cobra | Bile duct adenoma | Cowan, 1968 |
| *Naja nigricollis* | Black-necked Spitting Cobra | Osteochondrosarcoma (Metastatic to Spleen and Kidney) | Machotka and Jacobson, 1989 |
| *Naja nivea* ♂ | Cape Cobra | Bronchogenic adenocarcinoma | Effron et al., 1977 |
| *Naja sp.* (93-SnN-82) | Cobra | Hepatoma, malignant | Frye, 1994 |
| *Naja sp.* (haje?) | Egyptian cobra | Lymphosarcoma (heart and liver) | Cowan, 1968 |
| *Ophiophagus hannah* | King Cobra | Lymphoma | Willette et al., 2001 |
| *Ophiophagus hannah* ♀ | King Cobra | Tubular adenoma (kidney) | Catão-Dias and Nichols, 1999 |
| *Ophiophagus hannah* ♀ | King Cobra | Chondrosarcoma (vertebra) | Vico et al., 2016 |
| *Pseudechis porphyriacus* ♀ | Red-bellied Black Snake | Papillomas of skin; adenomatous proliferation of intrahepatic bile ducts | Effron et al., 1977 |
| *Pseudechis sp.* | Black Snake | Fibrosarcoma | Wadsworth, 1956 |
| *Walterinnesia aegyptia* | Sinai Desert Cobra | Renal adenocarcinoma | Frye, 1991a |
| *Walterinnesia sp.* (88-N-01) | Black Desert Cobra | Renal tubular adenoma | Frye, 1994 |
| “Elapid” | Cobra (?) | Adenocarcinoma (Pancreas) | Ratcliffe, 1943 |

<sup>1</sup>Possibly *Acrantophis madagascariensis*, which is not an Elapid.

**Table 2.**
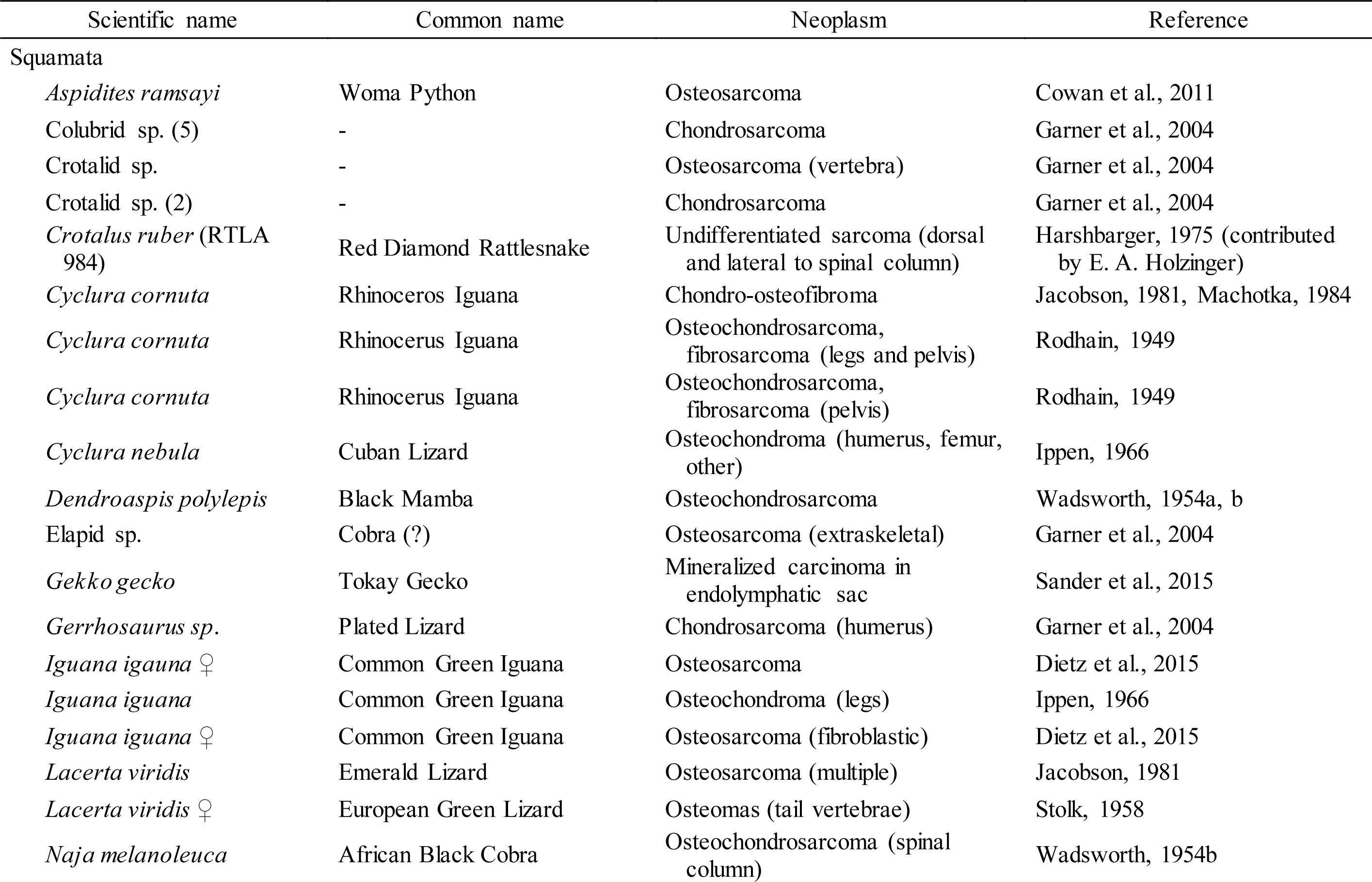
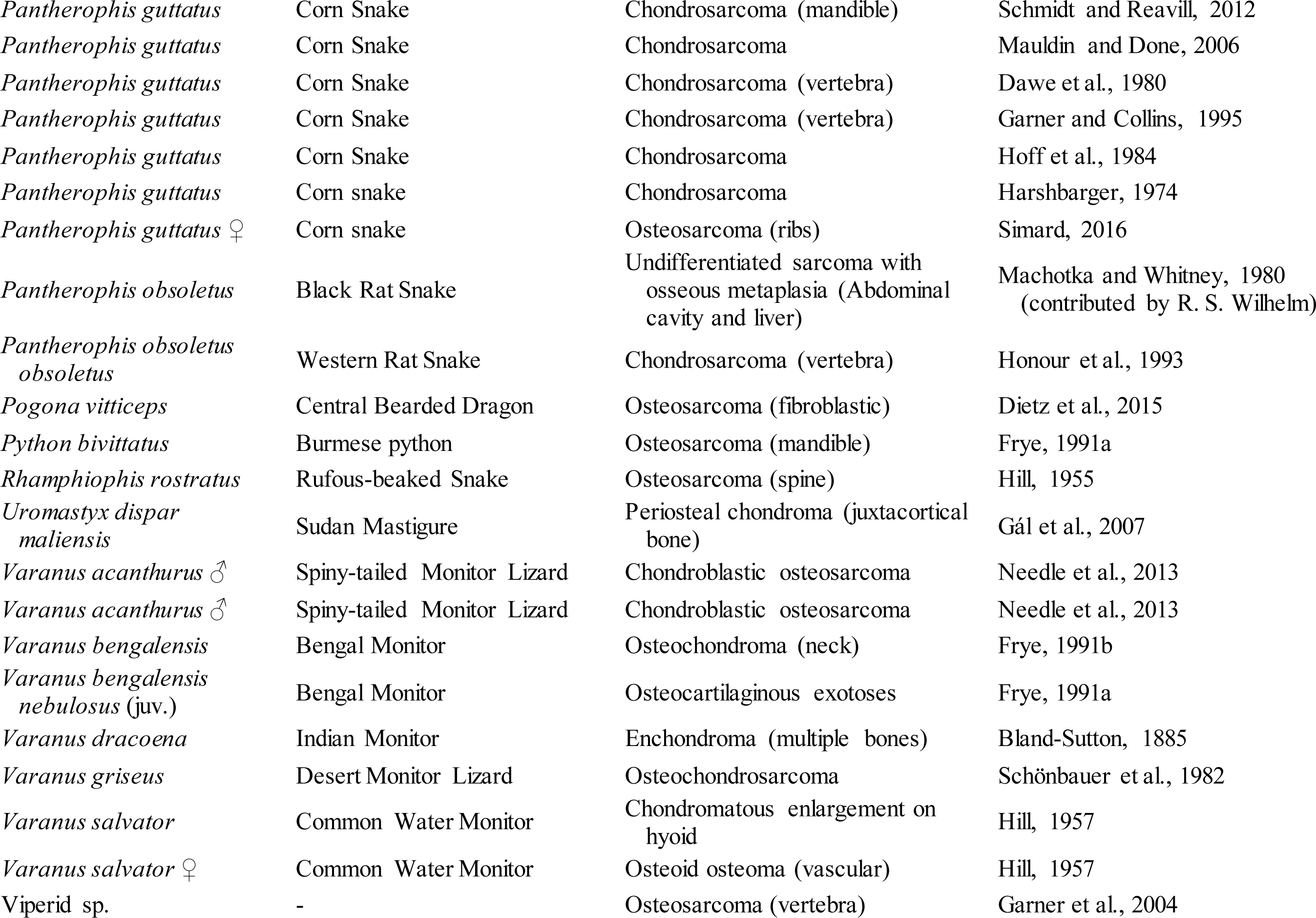
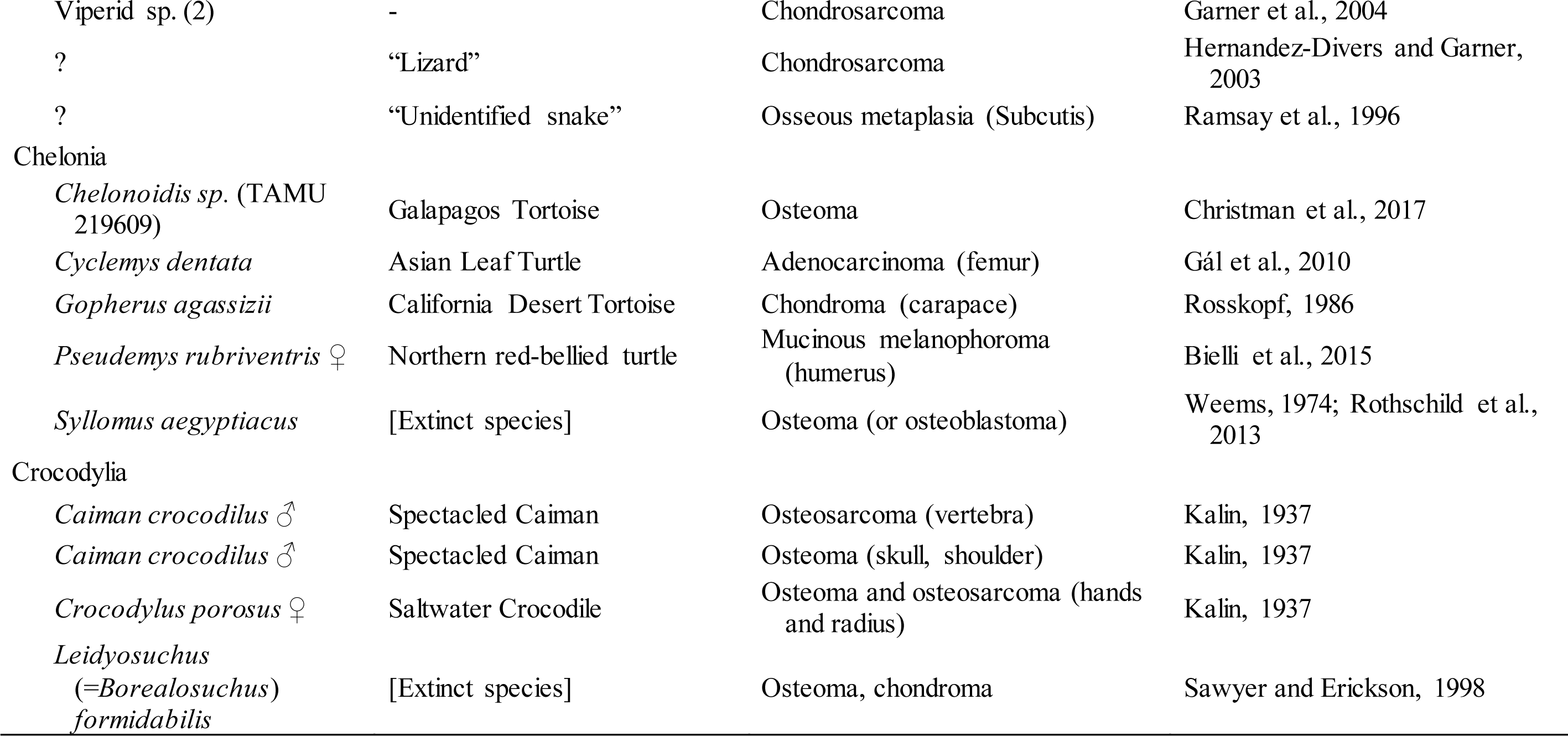
Summary of bony neosplasms reported in non-avian reptiles.

| Scientific name | Common name | Neoplasm | Reference |
| --- | --- | --- | --- |
| **Squamata** |  |  |  |
| *Aspidites ramsayi* | Woma Python | Osteosarcoma | Cowan et al., 2011 |
| Colubrid sp. (5) | - | Chondrosarcoma | Garner et al., 2004 |
| Crotalid sp. | - | Osteosarcoma (vertebra) | Garner et al., 2004 |
| Crotalid sp. (2) | - | Chondrosarcoma | Garner et al., 2004 |
| *Crotalus ruber* (RTLA 984) | Red Diamond Rattlesnake | Undifferentiated sarcoma (dorsal and lateral to spinal column) | Harshbarger, 1975 (contributed by E. A. Holzinger) |
| *Cyclura cornuta* | Rhinoceros Iguana | Chondro-osteofibroma | Jacobson, 1981, Machotka, 1984 |
| *Cyclura cornuta* | Rhinoceros Iguana | Osteochondrosarcoma, fibrosarcoma (legs and pelvis) | Rodhain, 1949 |
| *Cyclura cornuta* | Rhinoceros Iguana | Osteochondrosarcoma, fibrosarcoma (pelvis) | Rodhain, 1949 |
| *Cyclura nebula* | Cuban Lizard | Osteochondroma (humerus, femur, other) | Ippen, 1966 |
| *Dendroaspis polylepis* | Black Mamba | Osteochondrosarcoma | Wadsworth, 1954a, b |
| Elapid sp. | Cobra (?) | Osteosarcoma (extraskeletal) | Garner et al., 2004 |
| *Gekko gecko* | Tokay Gecko | Mineralized carcinoma in endolymphatic sac | Sander et al., 2015 |
| *Gerrhosaurus sp.* | Plated Lizard | Chondrosarcoma (humerus) | Garner et al., 2004 |
| *Iguana iguana* ♀ | Common Green Iguana | Osteosarcoma | Dietz et al., 2015 |
| *Iguana iguana* | Common Green Iguana | Osteochondroma (legs) | Ippen, 1966 |
| *Iguana iguana* ♀ | Common Green Iguana | Osteosarcoma (fibroblastic) | Dietz et al., 2015 |
| *Lacerta viridis* | Emerald Lizard | Osteosarcoma (multiple) | Jacobson, 1981 |
| *Lacerta viridis* ♀ | European Green Lizard | Osteomas (tail vertebrae) | Stolk, 1958 |
| *Naja melanoleuca* | African Black Cobra | Osteochondrosarcoma (spinal column) | Wadsworth, 1954b |
| *Pantherophis guttatus* | Corn Snake | Chondrosarcoma (mandible) | Schmidt and Reavill, 2012 |
| *Pantherophis guttatus* | Corn Snake | Chondrosarcoma | Mauldin and Done, 2006 |
| *Pantherophis guttatus* | Corn Snake | Chondrosarcoma (vertebra) | Dawe et al., 1980 |
| *Pantherophis guttatus* | Corn Snake | Chondrosarcoma (vertebra) | Garner and Collins, 1995 |
| *Pantherophis guttatus* | Corn Snake | Chondrosarcoma | Hoff et al., 1984 |
| *Pantherophis guttatus* | Corn snake | Chondrosarcoma | Harshbarger, 1974 |
| *Pantherophis guttatus* ♀ | Corn snake | Osteosarcoma (ribs) | Simard, 2016 |
| *Pantherophis obsoletus* | Black Rat Snake | Undifferentiated sarcoma with osseous metaplasia (Abdominal cavity and liver) | Machotka and Whitney, 1980 (contributed by R. S. Wilhelm) |
| *Pantherophis obsoletus obsoletus* | Western Rat Snake | Chondrosarcoma (vertebra) | Honour et al., 1993 |
| *Pogona vitticeps* | Central Bearded Dragon | Osteosarcoma (fibroblastic) | Dietz et al., 2015 |
| *Python bivittatus* | Burmese python | Osteosarcoma (mandible) | Frye, 1991a |
| *Rhamphiophis rostratus* | Rufous-beaked Snake | Osteosarcoma (spine) | Hill, 1955 |
| *Uromastix dispar maliensis* | Sudan Mastigure | Periosteal chondroma (juxtacortical bone) | Gál et al., 2007 |
| *Varanus acanthurus* ♂ | Spiny-tailed Monitor Lizard | Chondroblastic osteosarcoma | Needle et al., 2013 |
| *Varanus acanthurus* ♂ | Spiny-tailed Monitor Lizard | Chondroblastic osteosarcoma | Needle et al., 2013 |
| *Varanus bengalensis* | Bengal Monitor | Osteochondroma (neck) | Frye, 1991b |
| *Varanus bengalensis nebulosus* (juv.) | Bengal Monitor | Osteocartilaginous exostoses | Frye, 1991a |
| *Varanus dracoena* | Indian Monitor | Enchondroma (multiple bones) | Bland-Sutton, 1885 |
| *Varanus griseus* | Desert Monitor Lizard | Osteochondrosarcoma | Schönbauer et al., 1982 |
| *Varanus salvator* | Common Water Monitor | Chondromatous enlargement on hyoid | Hill, 1957 |
| *Varanus salvator* ♀ | Common Water Monitor | Osteoid osteoma (vascular) | Hill, 1957 |
| Viperid sp. | - | Osteosarcoma (vertebra) | Garner et al., 2004 |
| Viperid sp. (2) | - | Chondrosarcoma | Garner et al., 2004 |
| ? | “Lizard” | Chondrosarcoma | Hernandez-Divers and Garner, 2003 |
| ? | “Unidentified snake” | Osseous metaplasia (Subcutis) | Ramsay et al., 1996 |
| **Chelonia** |  |  |  |
| *Chelonoidis* sp. (TAMU 219609) | Galapagos Tortoise | Osteoma | Christman et al., 2017 |
| *Cyclenys dentata* | Asian Leaf Turtle | Adenocarcinoma (femur) | Gál et al., 2010 |
| *Gopherus agassizii* | California Desert Tortoise | Chondroma (carapace) | Roskopf, 1986 |
| *Pseudemys rubriventris* ♀ | Northern red-bellied turtle | Mucinous melanophoroma (humerus) | Bielli et al., 2015 |
| *Syllomus aegyptiacus* | [Extinct species] | Osteoma (or osteoblastoma) | Weems, 1974; Rothschild et al., 2013 |
| **Crocodylia** |  |  |  |
| *Caiman crocodilus* ♂ | Spectacled Caiman | Osteosarcoma (vertebra) | Kalin, 1937 |
| *Caiman crocodilus* ♂ | Spectacled Caiman | Osteoma (skull, shoulder) | Kalin, 1937 |
| *Crocodylus porosus* ♀ | Saltwater Crocodile | Osteoma and osteosarcoma (hands and radius) | Kalin, 1937 |
| *Leidyosuchus*<br>(= *Borealosuchus*)<br>*formidabilis* | [Extinct species] | Osteoma, chondroma | Sawyer and Erickson, 1998 |

Currently, the RTLA resides at Experimental Pathology Laboratories in Virginia. Jeffrey Wolf, current curator of the RTLA, knew of only one bony tumor in the Elapidae from their collection: a metastatic vertebral osteochondrosarcoma in *Naja nigricolis* (RTLA case #2474). Wolf noted no tumors in the collection within the genus *Micrurus*. Additionally, Wolf reported the following elapid neoplasms in the RTLA archive: six renal adenocarcinomas, two hepatocarcinomas, two lymphomas, one fibrosarcoma, one nasal carcinoma, one leiomyosarcoma, one Sertoli cell tumor, and one adrenal adenoma.

## Discussion

There are few alternative explanations for the displayed osteopathy. One alternative is osteoperiostitis. There is a precedent for this condition in two female Black Iguanas *(Ctenosaura similis)* and a female Chinese Stripe-tailed Snake *(Elaphe taenura;* Cowan, 1968). In each case, the bone and adjacent soft tissue became indistinct, softened, and exhibited exotosis. The periosteum of the laminae was affected, but not the intervertebral or apophyseal joints. Microscopically, the periosteum separated the new growth from the original bone, as new growth arose from the mesenchymal tissue on the surface of the periosteum. Cowan (1968) compared these osteopathic growths to virally induced osteoperiostitis (i.e., osteopetrosis) in chickens (Sanger et al., 1966). Compared to osteoperiostitis, as reported by Cowan (1968), the specimen in this study had fused intervertebral (i.e., condyle and cotyle) and apophyseal joints. Additionally, the growth occurred, it seems, inside of existing bone and not on top of the periosteum.

A second alternative is the possibility of infection of the bone or its supporting tissues (Rothschild et al., 2012). These authors exhaustively review and in some cases re-evaluate osteopathological conditions reported in the herpetological literature and rule several cases to be due to bone infection (Rothschild et al., 2012). In this study, infection cannot be ruled out, but there is clear evidence for neoplastic growth: low anisotropy of the bone, fused bones, and absence of bony growth on and in the periosteum. Furthermore, the scarcity of this specimen led the respective curator to disallow destructive histopathology.

Tumor vasculature becomes interesting in this study since we could easily segment it from the specimen’s micro CT data (Figure 1D). Generally speaking, solid tumors are more vascularized than healthy tissues (Forster et al., 2017), but their chaotic growth results in chronic and acute hypoxia due to poor circulation (Siemann, 2011; Forster et al., 2017). Among the earliest comments on vasculature of bony tumors in reptiles is Hill’s (1957) report of a vascular osteoid osteoma in a Common Water Monitor *(Varanus salvator).* Hill (1957) briefly notes that reptile osteomas (at least in snakes) may be synonymous with exotoses found in human osteomas. Interestingly, Stolk (1958) reports several osteomas in the caudal vertebrae of a female European Green Lizard *(Lacerta viridis)* that exhibited normal vasculature. The discrepancy in vasculature may be due to a different origin for the neoplastic growth in the *Lacerta,* as Stolk (1958) hypothesized the tumors to have originated from the marrow of the caudal long bones. Fitting with the generally chaotic nature of tumor vasculature, Cowan et al. (2011) reported an osteosarcoma in a Woma Python *(Aspidites ramsayi)* that was characterized by disorderly arrangement of osseous trabeculae. They also noted that mesenchymal cells surrounded an eosinophilic osteoid matrix.

Looking backwards through time, bony tumors do occasionally show up in the reptile fossil record (Moodie, 1917, 1923). A well-known example is Moodie’s (1918) description of an osteoma in a mosasaur (see also Moodie, 1923). In this case, however, Rothschild et al. (2012) reassessed this paleontological osteopathy as a hamartoma (i.e., not considered neoplasia). Our review also includes bony neoplasms in an extinct turtle, *Syllomus aegyptiacus* (Weems, 1974), and an extinct crocodile, *Leidyosuchus formidabilis* (Sawyer and Erickson, 1998). The fossil record, obviously, is sparse and uneven across time, and the condition and taphonomy of fossils complicates their accurate osteopathological diagnosis (Moodie, 1917; Rothschild et al., 2012). Computed tomography, especially from a synchotron light source, may assist properly diagnosing suspected fossil neoplasms, as histology is typically ruled out.

Prior to this report, we are unaware of any published neoplasms in new world elapid snakes (Table 1), likely reflecting their general absence in private collections and zoos compared to other species (Keck et al., 2011). Large, charismatic venomous cobras of the old world are relatively common in zoos throughout the world (e.g., Ramsay et al., 1996; Catão-Dias and Nichols, 1999; Collis and Fenili, 2011) and would thus be expected to be necropsied more often (Table 1). Compared to the hundreds of tumors reported for other families of snakes (Garner et al. 2004), even the RTLA (similar to other collections and reports), likely reflects the historical bias against including elapid snakes in zoo and private collections (Keck et al., 2011; Wolf J, pers. comm.). The relative obscurity of new world elapids (Schmidt, 1936), especially in captivity (Collis and Fenili, 2011), thus requires focused effort investigating specimens in natural history collections to understand their neoplasms. Another obvious omission in the neoplasm literature is a comparative phylogenetic study of tumors across reptiles and birds. Machotka and Jacobson (1989) provided the most recent phylogenetically organized treatment of neoplasms in this group; however, this study was entirely descriptive. Furthermore, our knowledge of comparative phylogenetics (e.g., Freckleton et al., 2002) and the reptile phylogeny has since vastly improved (Reeder et al. 2015, Streicher and Wiens, 2017). A comparative phylogentic test of neoplasia prevalence across the Sauria would reveal unexpected neoplasm absences and focus future study of natural history collections on these groups. Our report demonstrates that studying neoplasms is sometimes possible without dissecting valuable, rare specimens.

## Acknowledgements

We thank M Loocke for allowing complimentary use of the SMX-100CT. We also thank J Wolf for investigating our questions about tumors in the RTLA. A Beta Phi chapter Phi Sigma research grant to ASH funded the CT reconstruction computer.

---

**Video S1**. Section of vertebrae surrounding a vertebral osteosarcoma in Micrurus ancoralis (UTA R-55945). Available from MorphoSource: http://www.morphosource.org/Detail/MediaDetail/Show/media_id/34110.

**Video S2**. Pair of normal vertebrae in Micrurus ancoralis (UTA R-55945). Blood vessels are shown in red and connective tissues are shown in blue. Available from MorphoSource: http://www.morphosource.org/Detail/MediaDetail/Show/media_id/34108.

**Video S3**. Fused pair of pathologic vertebrae in Micrurus ancoralis (UTA R-55945). Blood vessels are shown in red, chaotic vasculature is shown in green, and connective tissues are shown in blue. Note the blunted and nearly absent neural spine and indistinguishable nature of the two vertebrae. Available from MorphoSource: http://www.morphosource.org/Detail/MediaDetail/Show/media_id/34109.

